# Breaking the cycle: How targeting CD44v6/MET signaling disrupts colorectal cancer cell plasticity

**DOI:** 10.1101/2024.09.27.614761

**Authors:** S. J. Sonnentag, S. Goldbach, D. Hoch, G. Andrieux, J. Becker, H. Fromm, L. Munoz-Sagredo, Y. M. Heneka, D. Grimm, F. R. Greten, Veronique Orian-Rousseau

## Abstract

The concept of cancer stemness is undergoing rapid evolution, facilitated by technological advances allowing the tracking and elimination of stem cells *in vivo*. Following their identification in solid tumors, cancer stem cells were placed at the center of tumor initiation, growth and metastasis. However, increasing evidence suggests that stemness is a cellular state that can be acquired by differentiated cells in a process called plasticity. Here we show that CD44v6 may act as a molecular switch controlling plasticity of colorectal cancer cells. CD44v6/MET signaling controls the reappearance of Lgr5^+^ cells after ablation, *in vitro* and *in vivo* as demonstrated in tumor organoids derived from *Lgr5*^*DTR/eGFP*^ mice using a CD44v6 blocking peptide. CD44v6 also affects the YAP/TAZ signaling pathway, involved in the very first steps of plasticity. In view of the essential role of plasticity for the establishment of metastases, blocking CD44v6 signaling may represent a pivotal and promising therapy.

**Graphical Abstract:** 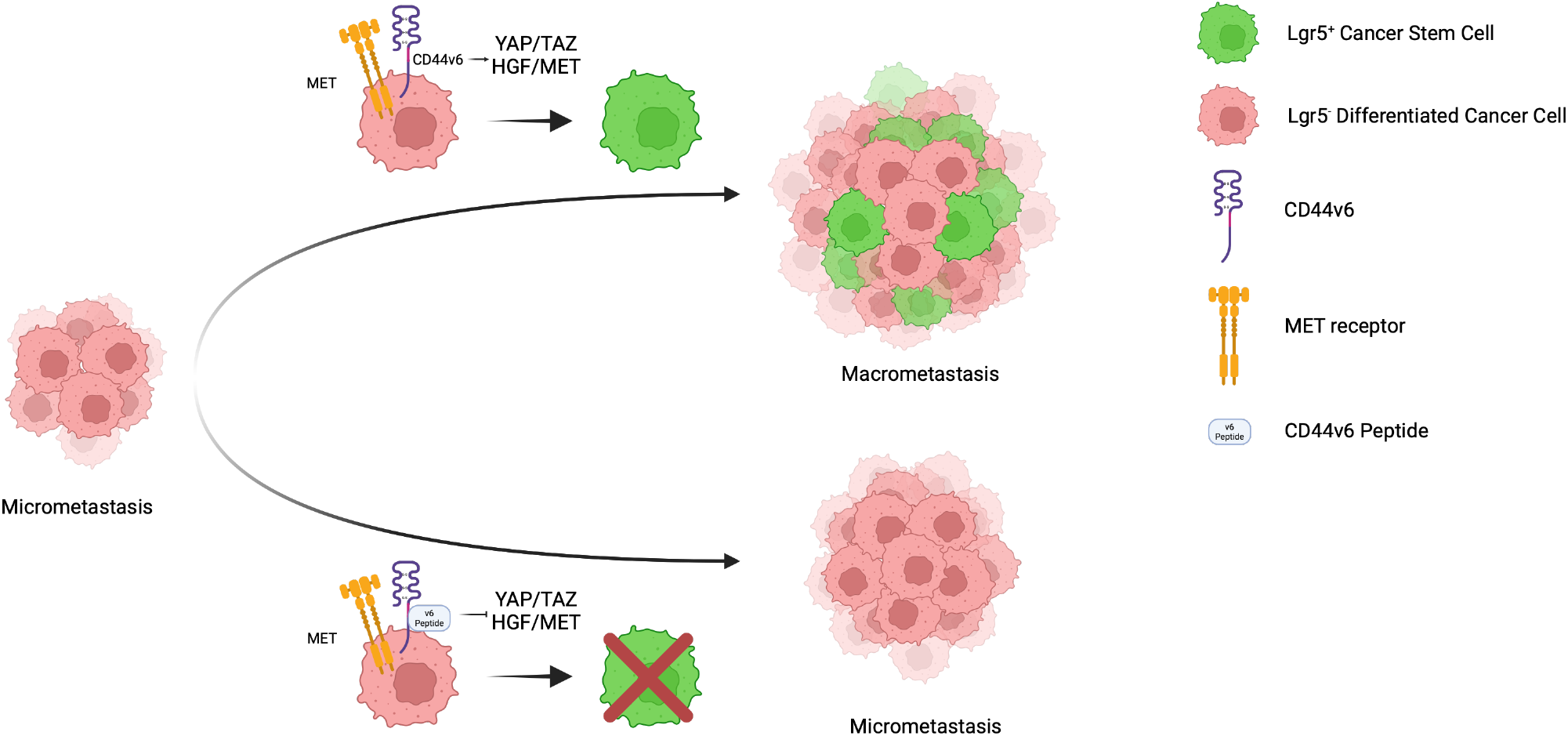

## Introduction

The term “cell plasticity” encompasses a range of mechanisms, including epithelial-mesenchymal transition (EMT) and its reciprocal process, mesenchymal-epithelial transition (MET), as well as trans-differentiation, referring to the transition of a cell from one lineage to another^1^. Although there is no consensus on what the term alludes to, its meaning has increasingly been used to describe the process of cell dedifferentiation from a differentiated state to that of a stem cell. In the intestine, stem cells (SCs) find themselves at the top of the cellular hierarchy. Lineage tracing experiments have shown that these SCs can give rise to all cells of the epithelium. Following the fate of SCs has been made possible with the engineering of the *Lgr5*_*eGFP*_ mouse model where eGFP is introduced into the gene locus of the *Lgr5* stem cell marker^2^. The addition of the diphtheria toxin receptor (DTR) sequence under the control of the same promoter (*Lgr5*^*DTR/eGFP*^ mice) allowed not only the tracking of Lgr5^+^ cells but also their ablation upon treatment with diphtheria toxin (DT)^3^.

These *Lgr5*^*DTR/eGFP*^ mice were crossed with *Apc*^*Min/+*^;*Kras*^*LSL-G12D*^;*Vil1*^*Cre*^ mice, designated as AKVL mice, allowing the tracking of Lgr5^+^ cells during colorectal tumor development and metastasis^4^. Using this and other models bearing additional mutations such as *p53* (AKVPL) and *Smad4* (AKVPSL), the group of F. de Sauvage showed that targeting Lgr5^+^ cells does not lead, as expected, to a shrinkage of a colorectal tumor but to tumor stasis. Following the depletion of Lgr5^+^ cells, the reappearance of these cells was observed one week after the discontinuation of DT treatment, highlighting the plasticity of Lgr5^-^ cells. Enrichment of Lgr5^+^ cells in micrometastases after the injection of AKVPSL organoids into mice suggested that dissemination and the initial steps of colonization are driven by Lgr5^+^ cells. However, challenging findings from the group of J. van Rheenen identified differentiated Lgr5^-^ cells as the most abundant disseminated cells in circulation5. The maintenance of the metastatic pool at distant locations was in turn dependent on the re-acquisition of stem cell properties. Indeed, upon arrival to the liver, these Lgr5^-^ cells underwent a dedifferentiation process into Lgr5^+^ cells mandatory for the establishment of metastasis.

The aforementioned findings have significant implications for cancer research. They indicate that the quest for drugs specifically targeting cancer stem cells may not yield the anticipated outcomes. The reappearance of cancer stem cells (CSC) from non-CSCs suggests that anti-CSC targeting drugs may prove ineffective. Conversely, targeting plasticity itself may prove a more beneficial approach.

In terms of pathways involved in the plasticity of colorectal cancer (CRC)-SCs, however, not a lot is known. Heinz et al.^6^ have proven the implication of the YAP/TAZ pathway in the first steps of plasticity. Decrease of the early-stage YAP activity was absolutely required for the growth of organoids and the establishment of cell-heterogeneity along with an increase of the Wnt signaling pathway.

The Wnt and YAP/TAZ signaling pathways have been associated with CD44, another important CSC marker^7^. The term CD44 designates a family of transmembrane glycoproteins that play a role in various physiological processes like proliferation, migration or survival^8^. The CD44 isoforms, generated through alternative splicing^7^, act as co-receptors for various other cell surface receptors including MET, VEGFR-2, EGFR or CXCR4^7^. CD44 is a member of the Wnt signalosome, in direct contact with the Wnt co-receptor LRP6^9^ and is involved in its maturation^10^. CD44, as a partner for LRP6, exerts a positive feedback loop necessary for intestinal regeneration^9^. In glioblastoma, CD44 is an upstream regulator of the mammalian Hippo pathway. Consequently, its downregulation results in the inactivation of YAP through phosphorylation^11^. Bringing this evidence together, CD44 emerges as a common denominator in the different signaling pathways involved in cell plasticity, and therefore an interesting target for CRC.

Plasticity of CRCs is enhanced by growth factors such as HGF^5^. Since CD44v6 is part of the MET signalosome controlling the activation and downstream signaling of the HGF/MET pair^12^, we also examine here the link between CD44v6, MET and plasticity.

In this paper, using publicly available single-cell RNA-seq (scRNA-seq) data obtained from samples of CRC cells (CRCCs) injected in the mesenteric vein and arriving at the liver, we show that CD44 isoforms are expressed at early stages of the plastic process. We demonstrate that plasticity of differentiated CRCCs depends on CD44/CD44v6 and as well on the interplay between CD44v6 and MET. CD44v6 also impacts the YAP/TAZ pathway, involved in the first steps of plasticity. The inhibition of CD44v6 by means of CD44v6 peptides decreases plasticity *in vivo*. Metastatic colorectal cancer is a major cause of mortality with a 5-year survival of 14%. Given the importance of the plastic process in metastasis of CRC, targeting this process through inhibition of the CD44/MET pair may prove to be decisive.

## Method Part

### Single-cell RNA-seq analysis

Transcript counts were downloaded from Gene Expression Omnibus (GSE189987) and processed using the Seurat R package (v4.3.2)^13^. Cells with fewer than 200 transcripts or more than 10% mitochondrial content were excluded from further analysis, following the protocol described by Heinz et al^6^. The filtered data were then processed as follows: data were log-normalized with a scale factor of 10,000 and scaled, while the percent mitochondrial content was regressed out. Principal Component Analysis (PCA) was performed on the top 5,000 most variable features. Louvain clustering was computed with a resolution of 0.8 on the nearest-neighbor graph (k=30) derived from the top 10 principal components. The identified clusters were visualized in two dimensions using UMAP. The over-representation of each sample (i.e., time point) was quantified with Fisher’s exact test. Marker gene analysis was conducted using a Wilcoxon test, comparing each cluster to all other clusters.

### Organoid maintenance

AKPL organoid were obtained from Jacco van Rheenen^5^ and cultured either in full murine intestinal growth medium (IntestiCult, STEMCELL Technologies Inc.) or under minimal medium conditions (Advanced DMEM/F12 (GibcoTM, Thermo Fisher Scientific), 4 mM Glutamax (GibcoTM, Thermo Fisher Scientific), B27 (2%; GibcoTM, Thermo Fisher Scientific), N-acetylcysteine (1.25 mM; Sigma) and Noggin (1%; PeproTech)). RMOs were cultured in murine intestinal growth medium (IntestiCult, STEMCELL Technologies Inc.)^14^.

### Flow cytometry

AKPL organoids and RMOs were removed from Matrigel (Corning) using Dispase (1 U/ml; STEMCELL Technologies Inc.) and mechanically dissociated using a 1 ml pipette tip. Afterwards, the remaining cell clusters were washed, incubated with TrypLE (GibcoTM, Thermo Fisher Scientific,) containing 4 µg/ml DNase I (Roche) for 10 min at 37 °C), filtered using a 70 µm strainer (Greiner-Bio) and resuspended in FACS-Buffer (PBS, 2% fetal bovine serum (GibcoTM, Thermo Fisher Scientific), 2 mM EDTA (Roth)). Flow cytometric analysis and sorting (FACS) was performed on a FACSAria™ Fusion (BD Biosciences) using the gating strategy depicted in Supplemental Figure 2. Data were analyzed using FlowJo software (BD Biosciences).

### Organoid formation assays

Lgr5^-^ cells from AKPL organoids and RMOs were obtained using the FACS strategy in Supplemental Figure 2. After sorting, 300 Lgr5^-^ cells per Matrigel (Corning) dome were seeded for observation and visual analysis of the plastic process. Developing organoids were counted manually using the Leica DM IL LED FLUO microscope (Leica) three and six days after seeding of FACS-sorted Lgr5^-^ or Lgr5^+^ cells into Matrigel (Corning).

### Inhibition assays

Lgr5^-^ cells from AKPL organoids and RMOs were obtained using the flow cytometry sorting strategy in Supplemental Figure 2. After sorting, 300 cells per Matrigel (Corning) dome were seeded for visual analysis of the plastic process. 3,000 cells were seeded for subsequent quantitative analysis, via quantitative polymerase chain reaction (qPCR) or FACS. To inhibit CD44, cells were treated with the panCD44 antibody (IM7) (100 µg/ml) or the corresponding IgG isotype control. To inhibit CD44v6, the species-specific mouse CD44v6 peptide (Intavis) (H-N***GWQ***G-OH)^15^ or the respective control peptide (Intavis), bearing 3 alanine instead of GWQ, was employed (100 nM). Direct MET inhibition was achieved using the MET/ALK inhibitor crizotinib (1 µM; Sigma-Aldrich). In all experimental conditions, the medium and the inhibitors were exchanged every two days for a maximum period of six days.

### Organoid size filtering and measurement

AKPL organoids were carefully removed from Matrigel (Corning) using a cut-off 1 ml pipette tip and Dispase (1 U/ml; STEMCELL Technologies Inc.) followed by centrifugation (300 x g for 5minutes at RT). Following aspiration of the supernatant, the organoids were resuspended in 1 ml Advanced DMEM/F12 (GibcoTM, Thermo Fisher Scientific) and applied to a pre-moistened 100 µm cell strainer (Greiner-Bio) positioned on top of a 50 ml non-adherent reaction tube. Organoids exceeding 100 µm in diameter were retained within the filter, subsequently recovered using a 1 ml pipette and 1 ml Advanced DMEM/F12, and finally transferred to a new 1.5 ml reaction tube. Organoids smaller than 100 µm in diameter or cells were able to pass the filter. To obtain organoids with a size of 40-70 µm and cells, the filtered suspension was filtered through a 70 µm cell strainer (Greiner-Bio) into another reaction tube, and subsequently through a 40 µm strainer (Greiner-Bio). The two fractions obtained (≧100 µm, 40-70 µm) were further processed for mRNA isolation and subsequent qPCR analysis. Measurement was performed manually using the Leica software tool and a Leica DM IL LED FLUO microscope (Leica).

### Immunofluorescent stainings

The staining of organoids was conducted on 8-well Ibidi microscopy slides (Ibidi). Following a three-days incubation period, the developed organoids were incubated with 4 µl of IM7-PE (BioLegend) (RMOs), IM7-AF700 (BioLegend) (AKPL organoids) or the respective directly coupled IgG for 30 minutes. Thereafter, the organoids were washed twice with PBS (GibcoTM, Thermo Fisher Scientific) and covered again with 250 µl of the respective medium. Three-dimensional images of the organoids were captured using the HC PL APO CS2 10x/0.40 DRY objective on a confocal microscope (Stellaris 5, Leica).

### Proliferation assay

Proliferation of organoids was analyzed using the Click-iT™ Plus EdU Alexa Fluor™ 647 Flowcytometry-Assay-Kit (Thermo Fischer Scientific). Lgr5^-^ cells were treated with the CD44v6-specific (100 nM, Intavis) or the corresponding control peptide (100 nM, Intavis) for a time period of six days. Afterwards, the organoids were carefully removed from the Matrigel (Corning) dome, dissociated into single cells, prepared according to the manufacturer’s protocol and analyzed by flow cytometry.

### Subcutaneous injection of AKPL organoids

10^5^ cells of dissociated AKPL organoids were resuspended as small cell clusters in IntestiCult organoid medium (STEMCELL Technologies Inc.) and Matrigel (Corning) (1:1), mixed to a final volume of 200 μl, and injected subcutaneously into the left flank of 8-12-weeks-old NOD.Cg-*Prkdc*^*scid*^*Il2rg*^*tm1Wjl*^/SzJ (*NSG*) mice (Jackson Laboratory). Mice were observed daily for the development of a subcutaneous tumor. Three weeks after injection, a tumor with an approximate volume of 400 mm^3^ had formed. To ablate Lgr5^+^ stem cells, diphtheria toxin (DT) (50 µg/kg in PBS; Sigma-Aldrich) was administered daily via intraperitoneal injection for seven days. Following elimination of the stem cells, the mice were injected *i*.*p*. with the CD44v6-specific peptide (1 µg/g in PBS; Intavis) or the AAA control peptide (1 µg/g in PBS; Intavis) three times a week for one week. On day eight after the last peptide injection, the animals were sacrificed by cervical dislocation, the tumors isolated, dissociated using the Tumor Dissociation Kit, mouse (Miltenyi Biotec), analyzed and sorted for the expression of Lgr5 using flow cytometry. All experiments were approved by the “Regierungspräsidium Karlsruhe” (35-9185/G-25/23). The animals were kept in the animal facility of the Karlsruhe Institute of Technology. Three animals were used per condition. Two control peptide mice had to be removed from the experiment after reaching the humane endpoint.

### mRNA isolation

Organoid mRNA was isolated using the RNeasy Mini Kit (Qiagen) according to the manufacturer’s protocol.

### cDNA preparation and pre-amplification

To obtain cDNA from freshly isolated mRNA, reverse transcription was performed using the qScriberTM cDNA Synthesis Kit from highQu according to the manufacturer’s protocol. Preamplification was included as a first step of cDNA sample processing due to the limited amount of starting material. Custom multiplex primer pools were set up consisting of gene-specific forward and reverse primers (see Table 1). *Cd44*-specific targeted preamplification was performed with a separate primer pool containing only two primers complementary to sequences in the 5’ and 3’ constant regions of the *Cd44*-mRNA (see Table 1). The cDNA samples were preamplified for 14 (in case of target gene expression) or 20 (*Cd44* analysis) cycles using SsoAdvanced PreAmp Supermix (Bio-Rad Laboratories) as directed by the manufacturer. Subsequently, the preamplification product was diluted in nuclease-free water for further use in downstream experiments.

### qPCR Analysis

Quantitative mRNA analysis was achieved using the CFX Connect Real-Time PCR Detection System (BioRad). Gene expression was normalized to the housekeeping gene *Gapdh* (Forward: ATGTGTCCGTCGTGGATCTGA; Reverse: TTGCTGTTGAAGTCGCAGGAG).

### *Cd44* exon-specific analysis

Qualitative *Cd44* exon analysis on the AKPL organoids and RMOs was carried out according to the PCR analysis of van Weering et al^16^ (see Table 2).

For quantitative analysis, a qPCR was performed using primers against the constant regions and for each variant exon. The resulting Cq values of the exons were put in relation to the Cq values of the constant regions.

### AAV transduction and evaluation

CT26 colorectal cancer cells were transfected with the STAR construct marking all Lgr5^+^ cancer stem cells by the expression of mNeonGreen^17^. To inhibit plasticity, these cells were first sorted for mNeonGreen^-^ cells and subsequently transduced with adeno-associated virus 2 (AAV2) vectors^18^ with a transduction ratio of 10^5^ vector genomes per cell. To downregulate CD44v6, two different shRNA-expressing vectors were used (sh35 and sh90). A scrambled shRNA-expressing vector served as control. Following transduction, the cells were immediately seeded in 50 µl hanging drops to form spheroids (1,000 cells per drop) and observed for the re-appearance of mNeonGreen. Analysis was performed on day three and day seven using a Zeiss LSM800 microscope. The re-appearance of mNeonGreen was calculated using the CTCF method: CTCF = (area * mean grey value) - (area * mean back background grey value)

## Results Part

### Expression of CD44 at early steps of plasticity

In order to place CD44 in the plasticity cascade, we re-analyzed publicly available scRNA-seq data sets from Heinz et al where a stepwise analysis of plasticity during CRC metastasis to the liver^6^ was made following injection of organoids into the circulation. Upon re-analysis, the transcriptional profiles were grouped in five clusters (0-4) spanning from day 1 to 27 (Figure 1A). An over-representation analysis highlights the enrichment of cells coming from day 1 within the cluster 0 (Figure 1B). Therefore, cluster 0 grouped genes are involved in the onset of the plastic process as for example YAP/TAZ^6^. *Cd44* is more highly represented in cluster 0 and barely expressed in cluster 1 and 2 (Figure 1C) suggesting a higher expression of *Cd44* in the first steps of plasticity. Additionally, *Cd44* is over-represented in cluster 3 (Figure 1C). This cluster represents late stages of metastatic outgrowth and is enriched in stem cell genes^6^. Given that CD44 is also involved in CRC stemness, its re-expression at late time points is expected.

**Figure 1:**
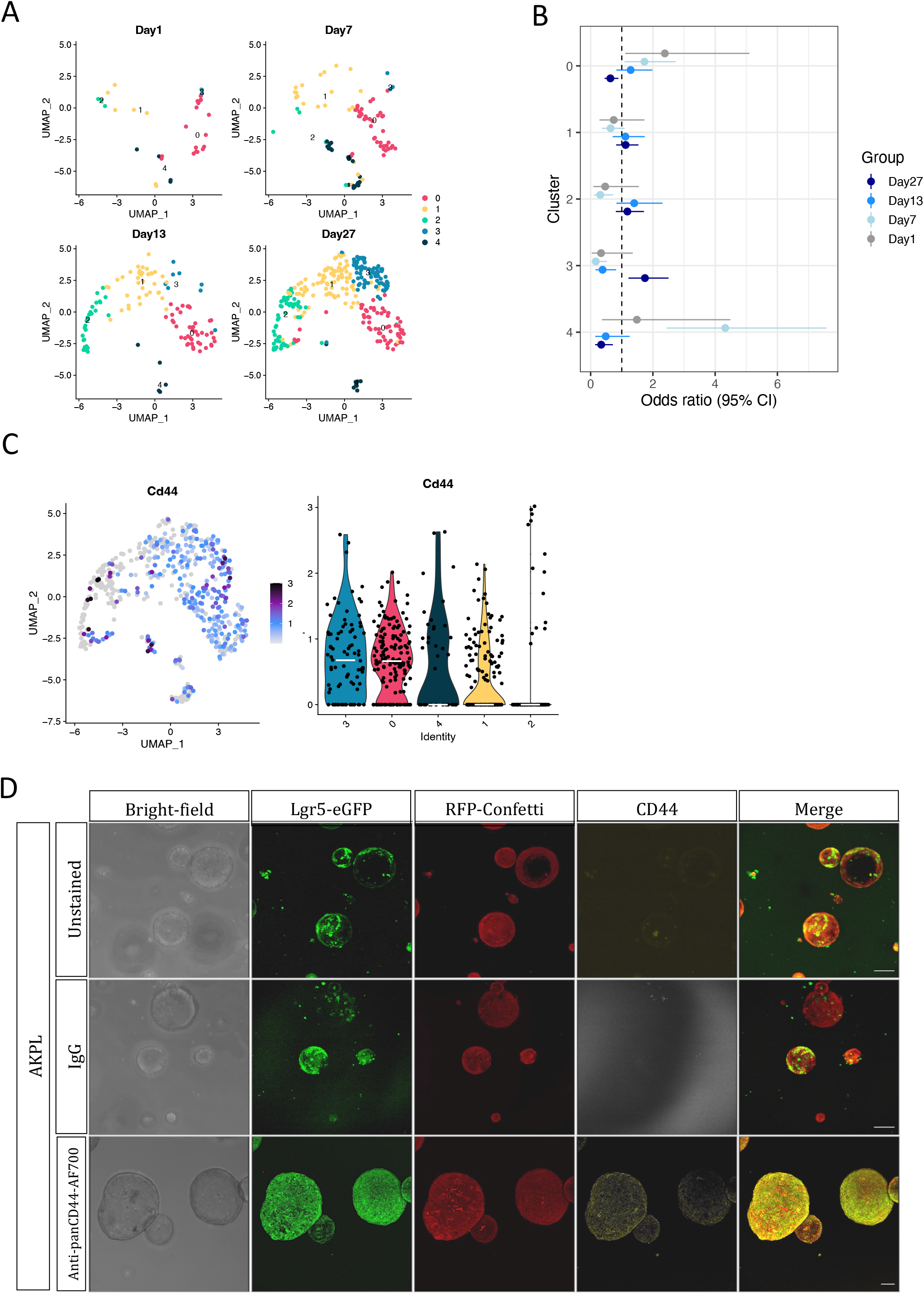
*Cd44* is expressed during the first steps of plasticity. A. UMAP visualization of the single-cell RNA-seq data. UMAP was split into four panels based on the time point (Day1, 7, 13 and 27). Louvain clusters are color-coded. B. Odds ratio plot from the over-representation analysis showing the enrichment of cells coming from specific days within each cluster. Error bars represent the 95% confidence interval. C. *Cd44* RNA expression visualized on the UMAP. Color scale represent the normalized expression. Violin plot showing the *Cd44* normalized expression across the cluster. Cluster are sorted according to the median expression (white bar). D. Immunofluorescence staining against CD44 (yellow) on AKPL organoids. Corresponding IgG and unstained served as controls. Scale bar: 100 µm

**Figure 2:**
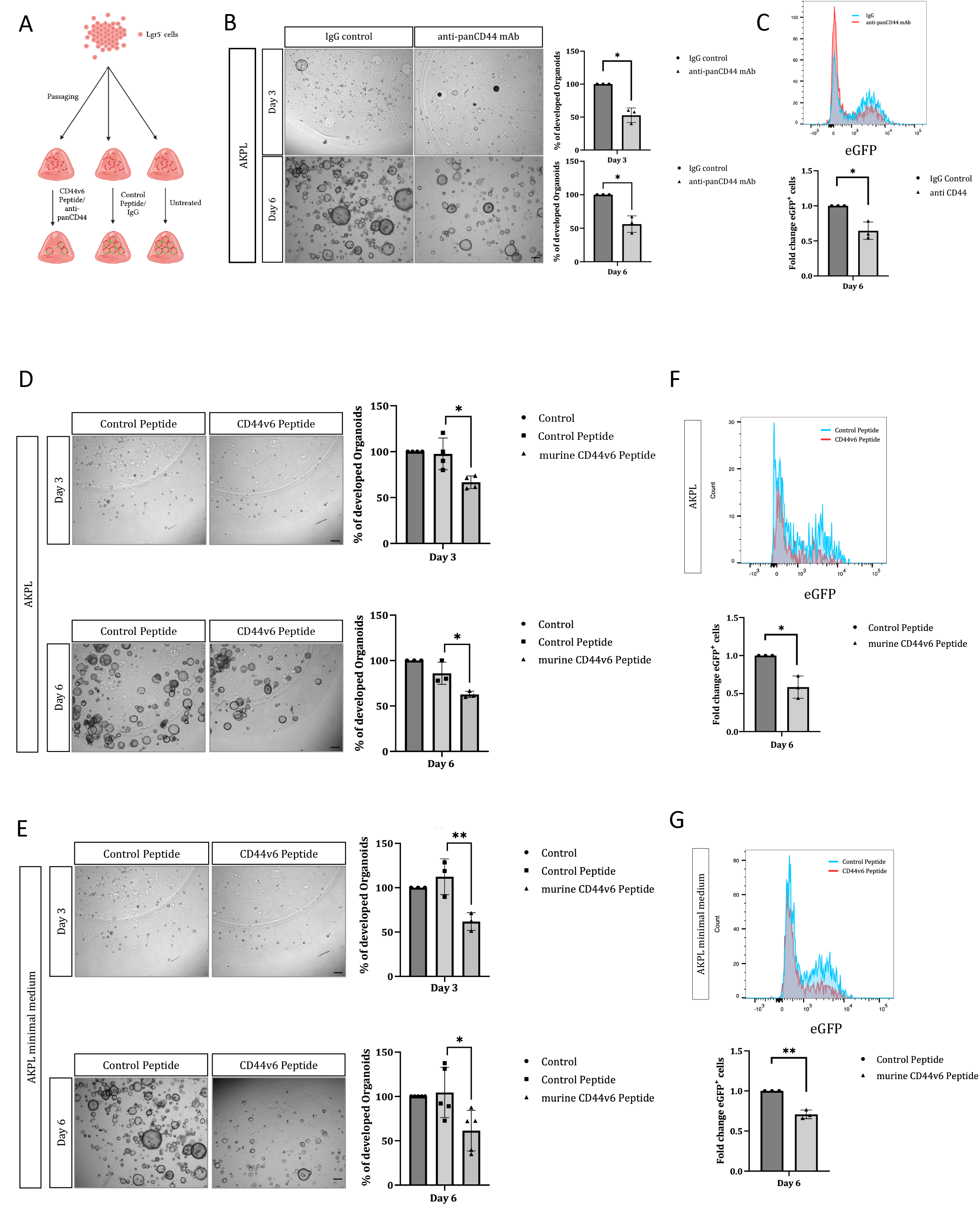
Inhibition of CD44 and CD44v6 impairs plasticity. A. Experimental procedure. Created in BioRender. Sonnentag, S. (2023) BioRender.com/h24u924 B. Blocking of CD44 using a panCD44 antibody (IM7) or IgG as control during the dedifferentiation of Lgr5^-^ into Lgr5^+^ AKPL cells (n=3 independent experiments) Values are presented as percentages *p-value<0.05. Representative pictures of organoids. Scale bar: 100 µm C. FC analysis six days after seeding of Lgr5^-^ cells cultured in full intestinal growth medium and treated with IM7 or IgG. Representative histogram and quantification of eGFP^+^ cells (n=3 independent experiments). Values are presented as fold changes. *p-value<0.05 D. Inhibition of CD44v6 by a CD44v6 peptide during the dedifferentiation process of Lgr5^-^ AKPL cells cultured in full intestinal growth medium. Representative pictures of day three and day six after seeding. Quantitative analysis of developed organoids (Day three: n=4 independent experiments; Day six: n=3 independent experiments). Values are presented as percentages. *p-value<0.05. Scale bar: 100 µm E. Inhibition of CD44v6 by a CD44v6 peptide during the dedifferentiation process of Lgr5^-^ AKPL cells cultured in minimal medium. Representative pictures of day three and day six after seeding. Quantitative analysis of developed organoids (Day three: n=3 independent experiments; Day six: n=5 independent experiments). Values are presented as percentages. *p-value<0.05, **p-value<0.01. Scale bar: 100 µm F. FC analysis of D. Representative histogram and quantification of the presence of eGFP at day six on AKPL cells treated with the CD44v6 or the respective control peptide (n=3 independent experiments). Values are presented as fold changes. *p-value<0.05 G. FC analysis of E. Representative histogram and quantification of eGFP at day six on AKPL cells treated with the CD44v6 or the respective control peptide (n=3 independent experiments). Values are presented as fold changes. **p-value<0.01

To further understand the potential role of CD44 in plasticity, we used organoids isolated from *Lgr5*^*DTR/eGFP*^ mice treated with AOM/DSS thereby inducing random mutations (RM) in the colon^14^. For simplification, these organoids will be designated RMO throughout the paper. We also used organoids isolated from *Lgr5*^*DTR/eGFP*^ mice crossed with the *VillinCreER*^*T2*^;*Apc*^*fl/fl*^;*Kras*^*LSL-g12D*^;*p53*^*KO/KO*^ and the *Confetti* mouse leading to CRC organoids harboring the classical mutations found in colorectal cancer^5^. These organoids were designated AKPL organoids as indicated in Fumagalli et al., 2020^5^.

AKPL organoids and RMOs express CD44 isoforms as shown by a yellow staining at the membrane of the organoids (Figure 1D and Supplemental Figure 1A, respectively). Run-off and relative exon quantification analysis revealed a lower expression of *Cd44s* (*Cd44* standard, smallest CD44 isoform) as compared to other *Cd44* isoforms including the longest variant isoform incorporating exons v1-v10 (Supplemental Figure 1B and C). Pan*Cd44* (all isoforms) and *Cd44v6* expression could be detected in both Lgr5^+^ and Lgr5^-^ cells isolated from AKPL organoids (Supplemental Figure 1D) in normal and minimal medium conditions and RMOs.

### Blocking CD44 inhibits cell plasticity

To examine the function of CD44 in plasticity, we isolated Lgr5^-^ cells via FACS and monitored the recurrence of Lgr5^+^ cells following the regrowth of AKPL organoids in the presence or absence of a CD44-blocking antibody directed against all CD44 isoforms (Figure 2A). The sorting and gating strategy for the Lgr5^-^ cells is shown in Supplemental Figure 2. At day three and day six (Figure 2B), the recovery of the Lgr5^+^ phenotype translated by the reappearance of cystic organoids was significantly decreased up to 50% upon incubation with the CD44 antibody (Figure 2B). Most importantly, the number of Lgr5^+^ (eGFP^+^) cells was decreased at day six upon inhibition of CD44 as shown by flow cytometry (FC) analysis in Figure 2C and the corresponding quantification. These data suggest that CD44 isoforms play an important role in the dedifferentiation of Lgr5-cells.

### The CD44v6/MET pair is involved in plasticity

Since the plastic process has been described to be boosted by growth factors such as HGF, we tested whether the co-receptor function of CD44v6 for MET^7^, the authentic receptor for HGF, was involved in this process. To do so, we made use of the CD44v6 peptide, a CD44v6-specific inhibitor, which blocks MET activation^15^. We isolated Lgr5^-^ cells from AKPL organoids and observed the plastic process during six days as described above. Treatment with the CD44v6 peptide reduced the generation of Lgr5^+^ from Lgr5^-^ cells by up to 30%, as measured by organoid reappearance, compared to treatment with the control peptide or PBS control (Figure 2D).

The same results were observed when plasticity of Lgr5^-^ cells was tested in minimal medium (Figure 2E). In contrast to full growth medium, the minimal medium does not contain additional niche factors like Wnt or EGF. These results suggest a cell-autonomous mechanism. We could not detect expression of HGF in AKPL organoids by qPCR (data not shown). However, this cannot be excluded since HGF secretion has been shown in early and late-stage CRC^19^.

The number of eGFP^+^ (Lgr5^+^) cells was decreased upon treatment of AKPL organoids with the CD44v6 peptide in both culture conditions (Figure 2F and 2G) as shown by FC. Although the CD44v6 peptide inhibited the generation of Lgr5^+^ cells from Lgr5^-^ in the case of the RMOs, a decrease in the number of stem cells was not observed (Supplemental Figure 3A and B).

**Figure 3:**
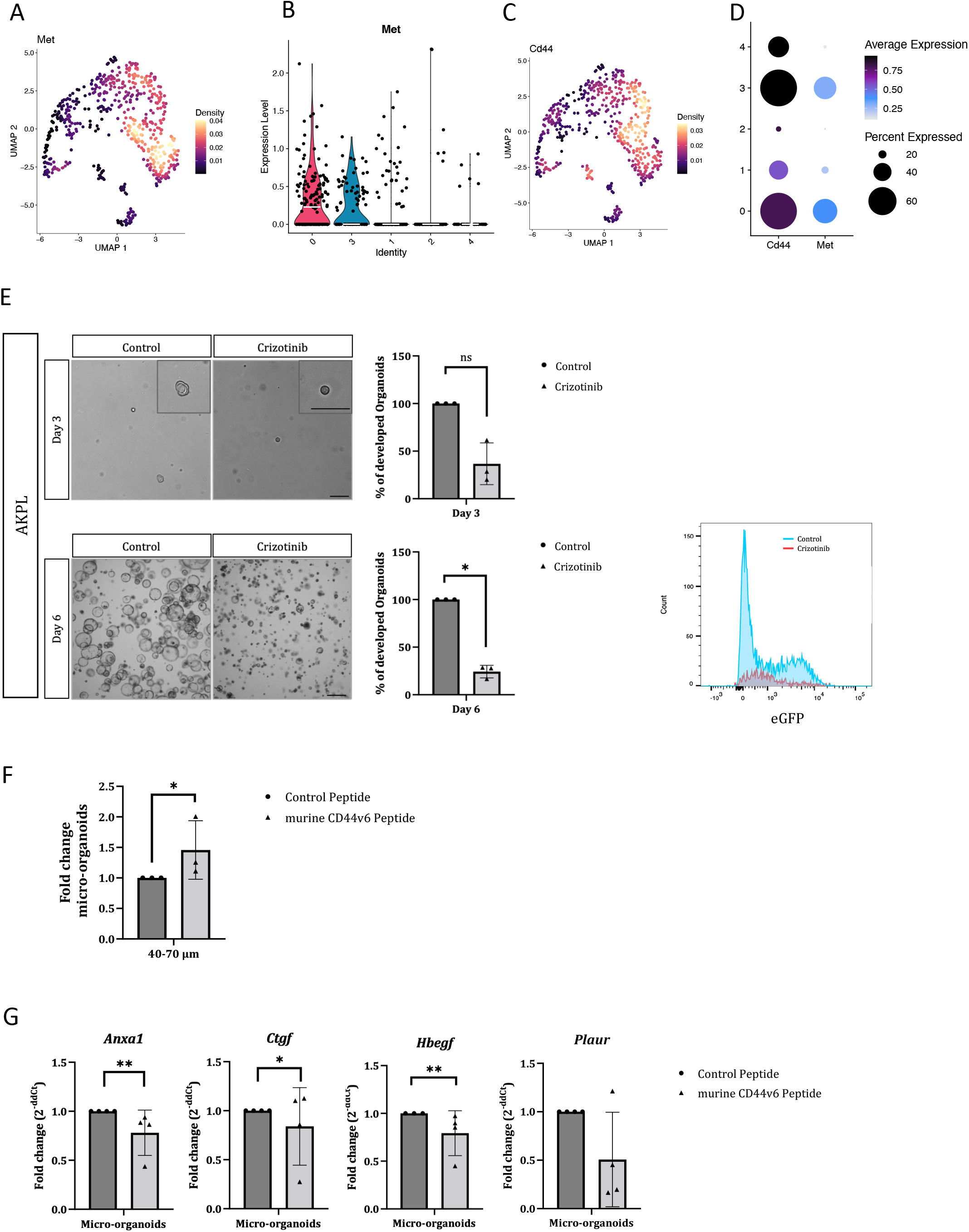
Influence of the CD44v6/MET axis on plasticity and YAP/TAZ signaling. A. Gene expression weighted 2D Kernel density plot of *Met* RNA expression visualized on the UMAP. B. Violin plot showing the *Met* normalized expression across the cluster. Clusters are sorted according to the median expression (white bar). C. Gene expression weighted 2D Kernel density plot of *Cd44* RNA expression visualized on the UMAP. D. Dotplot showing the percentage of cells expressing *Cd44* and *Met* (dot size) as well as the Average RNA expression of both genes across the Louvain clusters. E. Inhibition of MET/ALK during the dedifferentiation of Lgr5^-^ AKPL cells via crizotinib. Representative pictures of day three and day six after seeding. Quantitative analysis of developed organoids (Day three: n=3 independent experiments; Day six: n=5 independent experiments). Values are presented as percentages. *p-value<0.05, **p-value<0.01. Scale bar: 100 µm. Representative histogram of eGFP re-expression after plasticity upon treatment with crizotinib. F. Analysis of 40-70 µm micro-organoids during plasticity in AKPL organoids cultured in full intestinal growth medium and treated with the CD44v6 or control peptide for three days (n=3 independent experiments). Values are presented as fold-changes. *p-value<0.05 G. qPCR analysis of YAP/TAZ target genes in micro-organoids after plasticity (n=4 independent experiments). Values are presented as fold changes. *p-value<0.05, **p-value< 0.01

In order to distinguish an impact on dedifferentiation from an inhibition of proliferation, we performed an EdU-based proliferation assay in the presence and absence of the CD44v6 peptide during plasticity. The proliferation of the AKPL organoids was not affected by the treatment with the CD44v6 peptide in full growth (Supplemental Figure 3C) or minimal medium (Supplemental Figure 3D).

In conclusion, the data demonstrate that the inhibition of CD44v6 during the plasticity of Lgr5^-^ cells isolated from AKPL organoids results in a reduction of the number of stem cells, while the proliferation of cells remains unaltered. This evidence supports the hypothesis that the plasticity of Lgr5^-^ to become Lgr5^+^ cells is boosted by CD44v6.

MET is overexpressed in CRC^20,21^. Since CD44v6 directly interacts with MET^22^, we checked the expression profile of *Met* across the five clusters identified in the scRNA-seq dataset. Our analysis revealed that *Met* is actually over-expressed at the very first steps of the plastic process at the same time point (cluster 0) as *Cd44* (Figure 3A to C). Interestingly, *Met* and *Cd44* are co-expressed in cluster 0 (i.e. early time point) and cluster 3 as shown in Figure 3D. *Met* is expressed in Lgr5^+^ and Lgr5^-^ cells in a pattern similar to *Cd44* and *Cd44v6* (Supplemental Figure 1D).

We then directly inhibited MET using crizotinib, a MET/ALK inhibitor^23^. Lgr5^-^ cells were isolated from AKPL organoids and the recovery of the Lgr5^+^ expression was measured by organoid development. Crizotinib inhibited the plastic process to 80% (Figure 3E).

From these data, we conclude that the CD44v6/MET pathway is necessary for the induction of CRC plasticity. Both a direct targeting of MET or an indirect one through its association with CD44v6 has an impact on plasticity.

### A CD44/ YAP/TAZ collaboration is involved in the plastic process

The YAP/TAZ pathway, which is activated in micrometastases^6^, must be downregulated to allow the symmetry break and subsequent generation of cell-type heterogeneity that is essential for organoid and metastatic growth. Following plasticity of AKPL organoids in the presence of CD44v6 peptides, we observed an increased number of micro-organoids (defined by the size of 40-70 µm) (Figure 3F) accompanied by a decrease of macro-organoids (Figure 2D and E). This data suggests that blocking CD44v6 freezes the organoids in the micro-organoid state which then cannot undergo growth. At these very early steps of plasticity, we collected RNA from the organoids after FACS-based CSCs ablation. Expression of *Anxa1, Ctgf, Hbegf* and *Plaur*, all YAP/TAZ target genes, were decreased upon inhibition of CD44 (Figure 3G).

### In vivo inhibition of the plastic process by means of the CD44v6 peptide

In order to test the influence of CD44v6 on the plastic process *in vivo*, we injected the AKPL organoids subcutaneously in *NSG* mice, treated the mice with DT and then injected CD44v6 peptide or a control peptide intraperitoneally (Figure 4A). The FC analysis of the collected tumors revealed a significant decrease in the number of eGFP^+^ cells suggesting an impaired generation of Lgr5^+^ cells from Lgr5^-^ cells (Figure 4B). These results suggest that plasticity could be blocked by the CD44v6 peptide *in vivo*.

**Figure 4:**
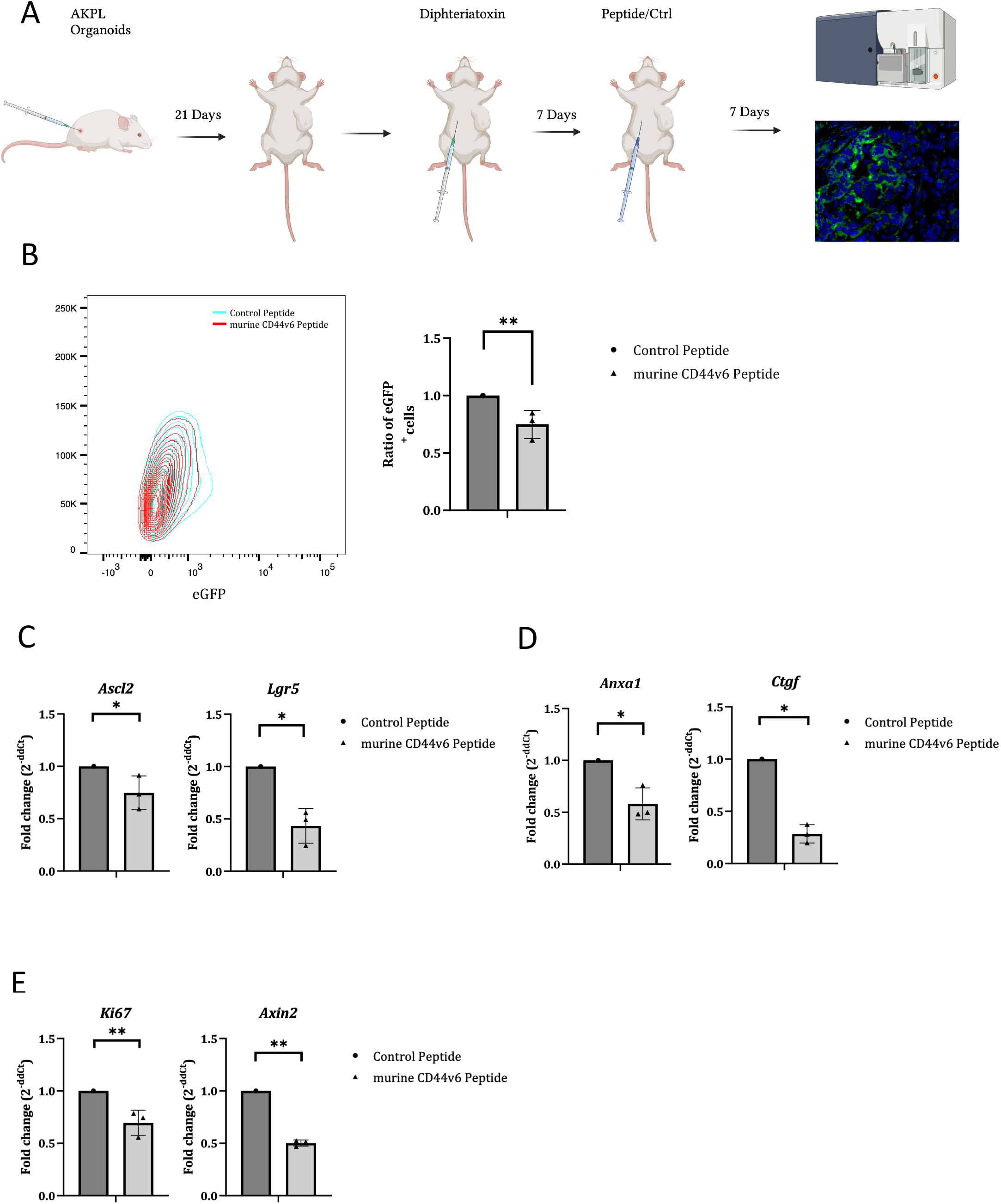
Inhibition of CD44v6 impairs plasticity *in vivo*. A. Experimental procedure. Created in BioRender. Sonnentag, S. (2023) BioRender.com/q06i684 B. FC analysis of the subcutaneous tumors after the ablation of Lgr5^+^ cells by DT treatment and subsequent inhibition of CD44v6 using the CD44v6-specific or control peptide (control peptide: n=1, CD44v6 peptide: n=3). Values are presented as ratios compared to the control condition. **p-value<0.01 C. qPCR analysis of CSC marker genes from isolated RFP^+^ cancer cells (control peptide: n=1, CD44v6 peptide: n=3). Values are presented as fold change compared to the control condition. *p-value<0.05 D. qPCR analysis of YAP/TAZ target genes from isolated RFP^+^ cancer cells (control peptide: n=1, CD44v6 peptide: n=3). Values are presented as fold change compared to the control condition. *p-value<0.05 E. qPCR analysis of proliferation and Wnt signaling marker genes from isolated RFP^+^ cancer cells. (control peptide: n=1, CD44v6 peptide: n=3). Values are presented as fold change compared to the control condition. **p-value<0.01 D. Comparison of *Cd44, Cd44v6* and *Met* expression in Lgr5^+^ and Lgr5^-^ cells from AKPL, AKPL cultures in minimal medium and RMOs (n=3 independent experiments). Values are presented as fold changes.

qPCR analysis of the red fluorescent protein (RFP^+^) marked cancer cells revealed a lower expression of cancer stem cell markers, including *Lgr5* and *Ascl2* (Figure 4C), in addition to a reduction in the expression of YAP/TAZ target genes (Figure 4D).

Of note, qPCR analysis of the RFP^+^ tumor cells also revealed a decrease in *Ki67* expression suggesting a reduced proliferation *in vivo* (Figure 4E). The Wnt target gene *Axin2* is as well downregulated.

## Discussion

In this paper, we show that the CD44v6/MET axis plays an important role during the process of plasticity of CRCCs. Using Lgr5^-^ cells isolated from CRC organoids, we were able to diminish the reappearance of Lgr5^+^ cells by means of a CD44v6 peptide. In addition, CD44v6 inhibition also significantly reduced the expression of YAP/TAZ target genes. Plasticity of CRCCs from AKPL organoids examined *in vivo* upon subcutaneous injection in *NSG* mice was also inhibited by the CD44v6 peptide.

The study by Fumagalli et al. demonstrates that metastases are seeded by Lgr5^-^ cells, which subsequently undergo a dedifferentiation into Lgr5^+^ CRCCs to sustain metastasis^5^. Furthermore, Heinz et al. implicated the YAP/TAZ pathway in the first steps of plasticity and thereby a crucial role in the development of metastases^6^.

The analysis of the scRNA-seq data from this study^6^ indicated that CD44 was upregulated during the initial stages of metastasis formation, suggesting that CD44 might act through YAP/TAZ signaling. The function of CD44 was tested by inhibiting all CD44 isoforms or CD44v6 isoforms specifically. CD44 had previously been demonstrated to impede the Hippo pathway, thereby inducing YAP/TAZ^11^. Several molecules may be responsible for the activation of YAP/TAZ through CD44. The PI3K/Akt pathway has been put forth as a potential activator of YAP/TAZ through merlin phosphorylation, and CD44 has also been demonstrated to regulate YAP through RhoA^24^. Given that the inhibition of CD44/CD44v6 resulted in alterations of the expression of YAP/TAZ target genes, we propose that CD44 isoforms exert a regulatory influence on the plastic process, at least in part, through their control of the YAP/TAZ pathway.

A reduction in YAP/TAZ signaling has been identified as a key factor in a process known as “symmetry break”, which is essential for the transition from a homogenous to a heterogenous organoid structure and the generation of stem cells. It can be hypothesized that the inhibition of CD44, which results in a decrease in YAP/TAZ, should induce plasticity. However, our data indicate a reduction in dedifferentiation upon CD44 blockade. This apparent contradiction may indicate that the inhibition of CD44 has broader implications that extend beyond the inhibition of YAP/TAZ signaling alone. An alternative hypothesis is that the inhibition of CD44 prevents the threshold level of YAP/TAZ signaling required for the induction of plasticity.

Plasticity of differentiated Lgr5^-^ cells can be divided into distinct steps, including proliferation, acquisition of heterogeneity, dedifferentiation and subsequent proliferation, ultimately resulting in the formation of cystic organoids or established macrometastases comprising Lgr5^-^ and Lgr5^+^ cancer cells *in vivo*. Following treatment with the CD44v6 peptide, a notable reduction in the eGFP signal was observed, accompanied by an increase in the proportion of micro-organoids in the AKPL organoids. This was not observed for the RMOs, although the number of cystic organoids was decreased upon CD44v6 inhibition. One possible explanation for this discrepancy is that in the case of RMOs, disturbance of other important steps leads to the same net outcome.

Here, we also show a clear impact of the MET/CD44v6 pair on the plastic process. Active HGF/MET signaling has previously been shown to induce nuclear translocation of YAP resulting in the expression of YAP/TAZ target genes in pancreatic cancer cell lines^25^. The inhibition of MET at the membrane by means of the CD44v6 peptide might have a negative effect on YAP/TAZ signaling. The influence of MET on plasticity might also be linked to its ability to induce the PI3K pathway^26^ and consequently YAP.

It is also possible that the function of MET is independent of YAP/TAZ. MET and CD44v6 have been identified as key players in CRC metastasis. Their presence on CR-CSC has been demonstrated by Todaro et al. to be indispensable for the metastatic process^27^. Here, CD44v6/MET expression on Lgr5^-^ cells appears to be a prerequisite for their dedifferentiation into stem cells and subsequent metastasis. Although these observations may initially appear to be in conflict, they can be reconciled by the fact that plasticity is required for metastases maintenance. It is clear that CD44 is a marker for cancer stem cells, and that CD44 isoforms are expressed on Lgr5^+^ cells. However, v6-containing isoforms and other CD44 isoforms can also be detected on Lgr5^-^ cells (Supplemental Figure 1D).

Collectively, our data indicate that CD44, and in particular CD44v6, play a pivotal role in the plasticity of CRC through its involvement with the YAP/TAZ and MET pathways (Graphical Abstract). CD44v6 has been demonstrated to be involved in colorectal cancer initiation. Indeed, CD44v6 shRNA reduced the adenoma number in *Apc*^*Min/+*^ mice^28^. CD44v6 was also implicated in CRC metastasis, in conjunction with MET^27^. There, CD44v6 was linked to its ability to maintain the stem cell phenotype^27^. Altogether, CD44v6 emerges as a potential therapeutic target along the tumor/metastasis route. Therefore, we used adeno-associated virus vectors (AAVs) expressing an anti-CD44v6 shRNA on spheroids from the CT26 CRC cell line. These vectors drastically blocked the plastic process from Lgr5^-^ spheroids as shown in Supplemental Figure 4A and B, suggesting a potential therapeutic benefit and motivating their further preclinical assessment.

## Supporting information

Licence Biorender

Licence Biorender

Licence Biorender

## Acknowledgment

We acknowledge funding from the German Federal Ministry of Education and Research (BMBF) within the Medical Informatics Funding Scheme (EkoEstMed–FKZ 01ZZ2015 to GA). S.J.S. and V.O-R. were supported by the Helmholtz program “Materials Systems Engineering (MSE)”. V.O-R and D.G. thank the HeiKa Research Bridges Grant for funding. We thank the animal facility of IBCS-FMS. We thank Genentech for providing the *Lgr5*^*DTR/eGFP*^ mouse (MTA: OM-220494). We thank J. Van Rheenen (Netherlands Cancer Institute (NKI)) for providing the AKPL organoids (MTA: OM-220494 and D1DF0921-BECA-420F-970E-3Ec90524F00B) and Hugo Snippert (University Medical Center Utrecht) for giving us the STAR construct ^17^.

## Figure Legends

Graphical Abstract: CD44v6 signaling in colorectal cancer cell plasticity. Created in BioRender. Sonnentag, S. (2024) BioRender.com/v47g193

Supplemental Figure 1

A. Immunofluorescence staining of CD44 (yellow) on RMOs. Scale bar: 100 µm

B. Representative runoff analysis of *Cd44* isoform expression on AKPL organoids, AKPL cultured in minimal medium and RMOs.

C. Quantitative analysis of the expression of different *Cd44* isoforms from AKPL organoids, AKPL cultured in minimal medium and RMOs (n=3 independent experiments). Values are presented as percentages.

Supplemental Figure 2

Representative cytometric plots depicting the gating strategy for analysis and sorting of Lgr5^+^ and Lgr5^-^ cells of AKPL organoids in normal and minimal medium.

Supplemental Figure 3

A. Inhibition of CD44v6 during the dedifferentiation process of Lgr5^-^ RMO cells. Representative pictures of day three and day six after seeding. Quantitative analysis of developed organoids (Day three: n=4 independent experiments; Day six: n=4 independent experiments). Values are presented as percentages. ***p-value<0.001. Scale bar: 100 µm

B. FC analysis of eGFP re-expression upon inhibition of CD44v6 during plasticity in RMO cells. Representative histogram and quantification of eGFP at day six on RMO cells treated with the CD44v6 or the respective control peptide (n=3 independent experiments). Values are presented as fold changes. ns=non-significant

C. EdU-based proliferation assay of AKPL organoids cultured in full intestinal growth medium at day six after seeding of Lgr5^-^ cells (treated or not with the CD44v6 or respective control peptide). Representative histogram of EdU measurements via flow cytometry. Quantitative analysis of EdU^+^ and EdU^-^ cells (n=5 independent experiments). ns=non-significant

D. EdU-based proliferation assay of AKPL organoids cultured in minimal medium at day six after seeding of Lgr5^-^ cells (not treated or treated with CD44v6 or respective control peptides). Representative histogram of EdU measurements via flow cytometry. Quantitative analysis of EdU^+^ and EdU^-^ cells (n=4 independent experiments). ns=non-significant

Supplemental Figure 4

A. Representative images of CT26 spheroids seven days after infection with AAVs carrying shRNA Scramble or shRNA CD44v6 vectors. mNeonGreen: Lgr5^+^ cells. mTurquoise: infection efficiency. Scale bar: 100 µm

B. Quantitative analysis of the re-expression of mNeonGreen in CT26 cells. Values are presented as fold changes compared to day three of each condition (n=1 experiment using 5 different spheroids).

