## Supplementary material for "Breaking the cycle: How targeting CD44v6/MET signaling disrupts colorectal cancer cell plasticity": Licence Biorender

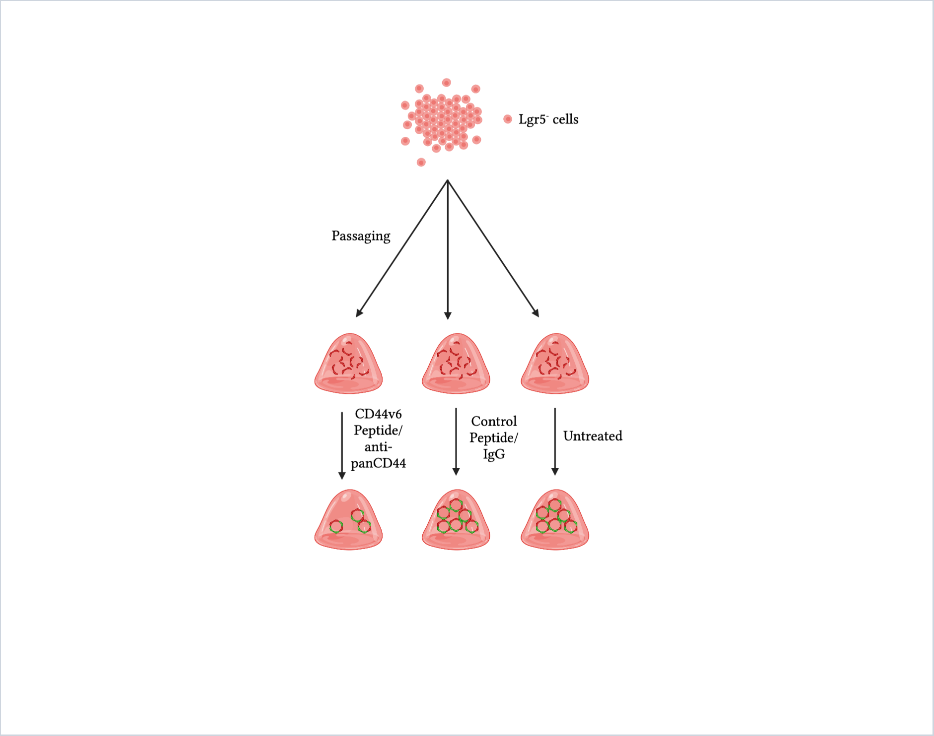


*For any questions regarding this document, or other questions about publishing with BioRender, please refer to our BioRender Publication Guide, or contact BioRender Support at.*
