## Supplementary material for "Breaking the cycle: How targeting CD44v6/MET signaling disrupts colorectal cancer cell plasticity": Table 1

Table 1: Primer sequences for RT-qPCR. The upper part of the table represents the primer sequences included in the multiplex preamplification primer pool.

| Primer name | sequence fwd primer 5'->3' | sequence rev primer 5'->3' |
|---|---|---|
| **Gapdh** | TCAACAGCAACTCCCACTCTTC | GGTGGTCCAGGGTTTCTTACTC |
| **Ascl2** | CTGCGAGGGAGAGCTAAGC | GCATAGGCCCAGGTTTCTTG |
| **Lgr5** | GTTGTCAGCCGTGGTCATAG | AGCCATTTGAGAGAGTGATGAAC |
| **Anxa1** | TCGCAGAGTGTTTCAGAATTACG | CACTTCTCAATGTCACCCTTCAG |
| **Ctgf** | CTGTGCCTGCCATTACAACTG | TCCCTTACTTCCTGGCTTTACG |
| **Ki67** | ACCCAGCACTCCAAAGAAACC | GTCGGGCAGGCACACTTAC |
| **Axin2** | GTGCAAACTCTCACCCACCGT | CGTCGCTGGATAACTCGCTGT |
| **Hbegf** | TCGTCCGTCTGTCTTCTTGTC | CACGCCCAACTTCACTTTCTC |
| **Plaur** | CTTCAGAGCTTTCCACCGAATG | GGTTTCCCAGCACATCTAAGC |
| **Met** | CACTGTCAAGGTTGCTGATTTCG | GGTGGTGAACTTCTGCGTTTG |
| **CD44c16c17 (pan)** | TCGAGACTCATCCAAGGACTCCA | CCACTGTCCTGGTTCGCACTT |
| **CD44s** | TCGAGAAGAGCACCCCAGAAAG | GTCTCGATCTCTGGTGGCCAAG |
| **CD44v1** | TGCCTCAACTGTGCACTCAAAA | ACTCGTTGTGGGCTCCTGAGT |
| **CD44v2** | GATGACCACCCCTGAAACACC | CAGGCATCTTCGTTGTTGTATGA |
| **CD44v3** | CGGAGTCAAATACCAACCCAAC | TCATCATCAATGCCTGATCCA |
| **CD44v4** | TGCAAGTACTCCACGGGTTTCTG | CCTGGTGGTTGTCTGGAGTAGT |
| **CD44v5** | ATATAGACAGAATCAGCACCA | CTCTTCATCCTGATACTCATG |
| **CD44v6** | AGCAGCTACCCAGCAGGAGAC | CCCTTCTGTCACATGGGAGTC |
| **CD44v7** | CGGCCCACAACAACCATCCAA | TTCTGTTTGATGACCTTGTCCCA |
| **CD44v8** | TCCAGTCATAGTACAACCCTTCAGC | CTGAAAGTGGTCCTGTCCTGTTCA |
| **CD44v9** | GGAGAGCCGGAAGAAGACGAA | GGCAGAATAGAAGTTGTTGGATGGT |
| **CD44v10** | GCGGCGCTAAAGATGCAAGAA | CCTCAGTTTTAGCAGGGGTCACT |
