## Supplementary material for "Breaking the cycle: How targeting CD44v6/MET signaling disrupts colorectal cancer cell plasticity": Table 2

Table 2: Primer sequences for Exon-specific PCR (aka. Run-Off-PCR) and the CD44-specific preamplification.

| Primer Name | Sequence 5'->3' | Remark |
|---|---|---|
| **CD44c16c17 rev** | CCACTGTCCTGGTTCGCACTT | = common reverse primer |
| **CD44 c3c4 fwd** | TCATCCCAACGCTATCTGTGCA | = CD44 pan amplification; CD44-specific preamplification |
| **CD44v1 fwd** | AAGCCATGCAGCAGCTCAGAA | |
| **CD44v2 fwd** | ACCCCTGAAACACCACCCAAGA | |
| **CD44v3 fwd** | CGGAGTCAAATACCAACCCAAC | |
| **CD44v4 fwd** | TGCAAGTACTCCACGGGTTTCTG | |
| **CD44v5 fwd** | CCGGAACCACAGCCTCCTTTC | |
| **CD44v6 fwd** | CAGCAGGAGACGTGGTTTCAG | |
| **CD44v7 fwd** | CGGCCCACAACAACCATCCAA | |
| **CD44v8 fwd** | TCCAGTCATAGTACAACCCTTCAGC | |
| **CD44v9 fwd** | GGAGAGCCGGAAGAAGACGAA | |
| **CD44v10 fwd** | GCGGCGCTAAAGATGCAAGAA | |
| **CD44s fwd** | CTTGGCCACCAGAGATCGAGAC | = CD44 standard amplicon |
