## Supplementary figures and images for "Breaking the cycle: How targeting CD44v6/MET signaling disrupts colorectal cancer cell plasticity"

### Supplemental Figures

#### Supplemental Figure 1

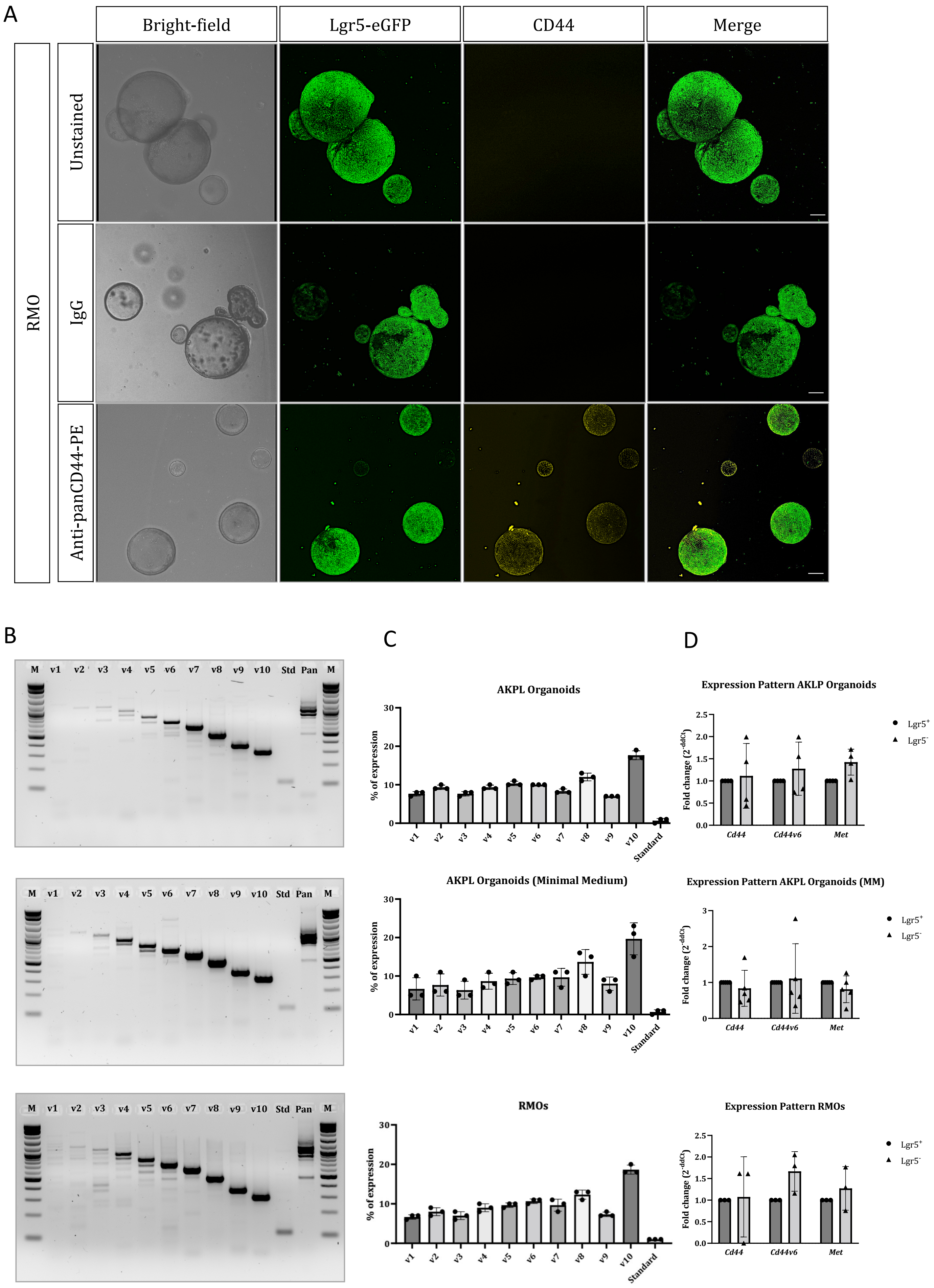

Supplemental Figure 2

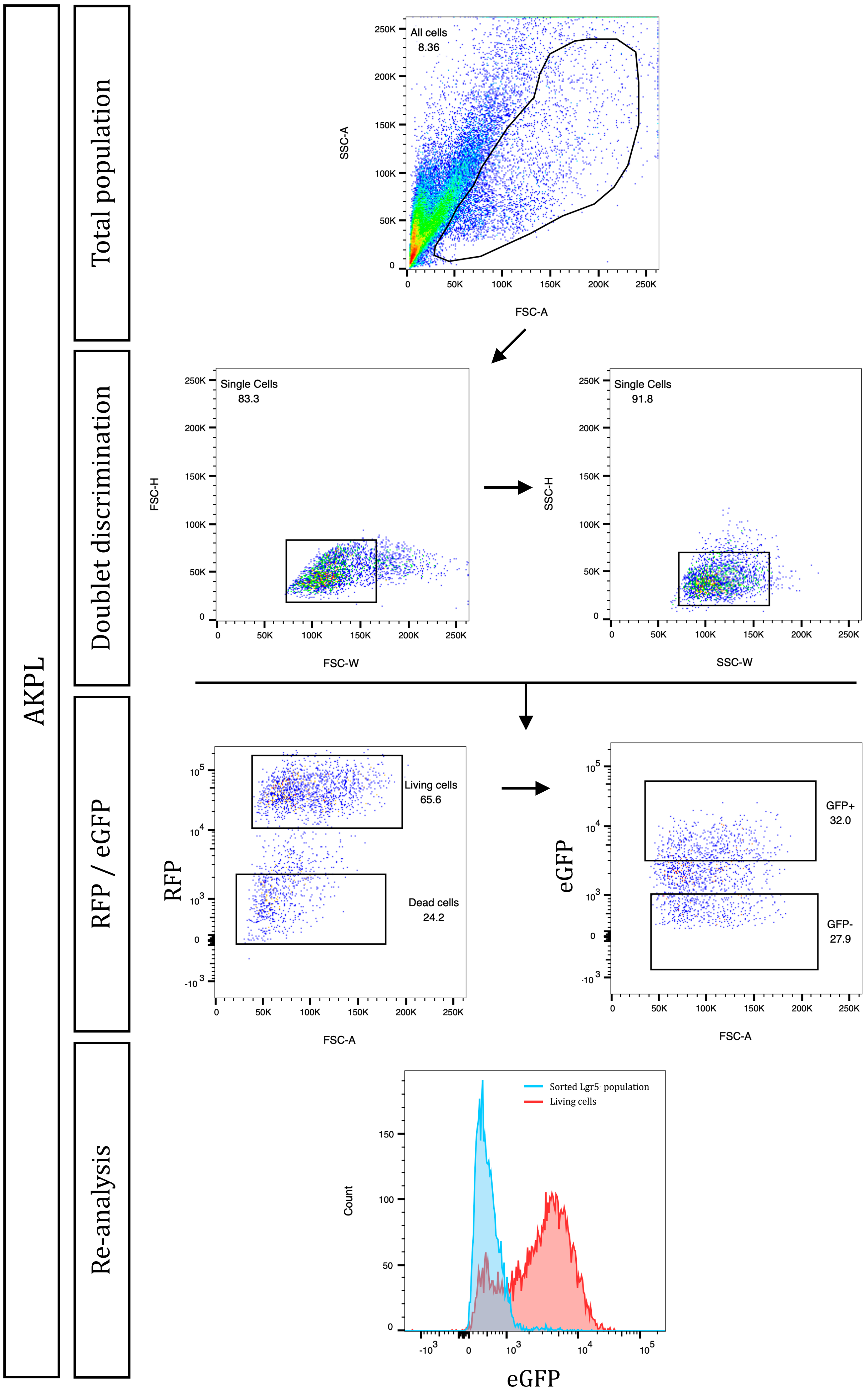

#### Supplemental Figure 3

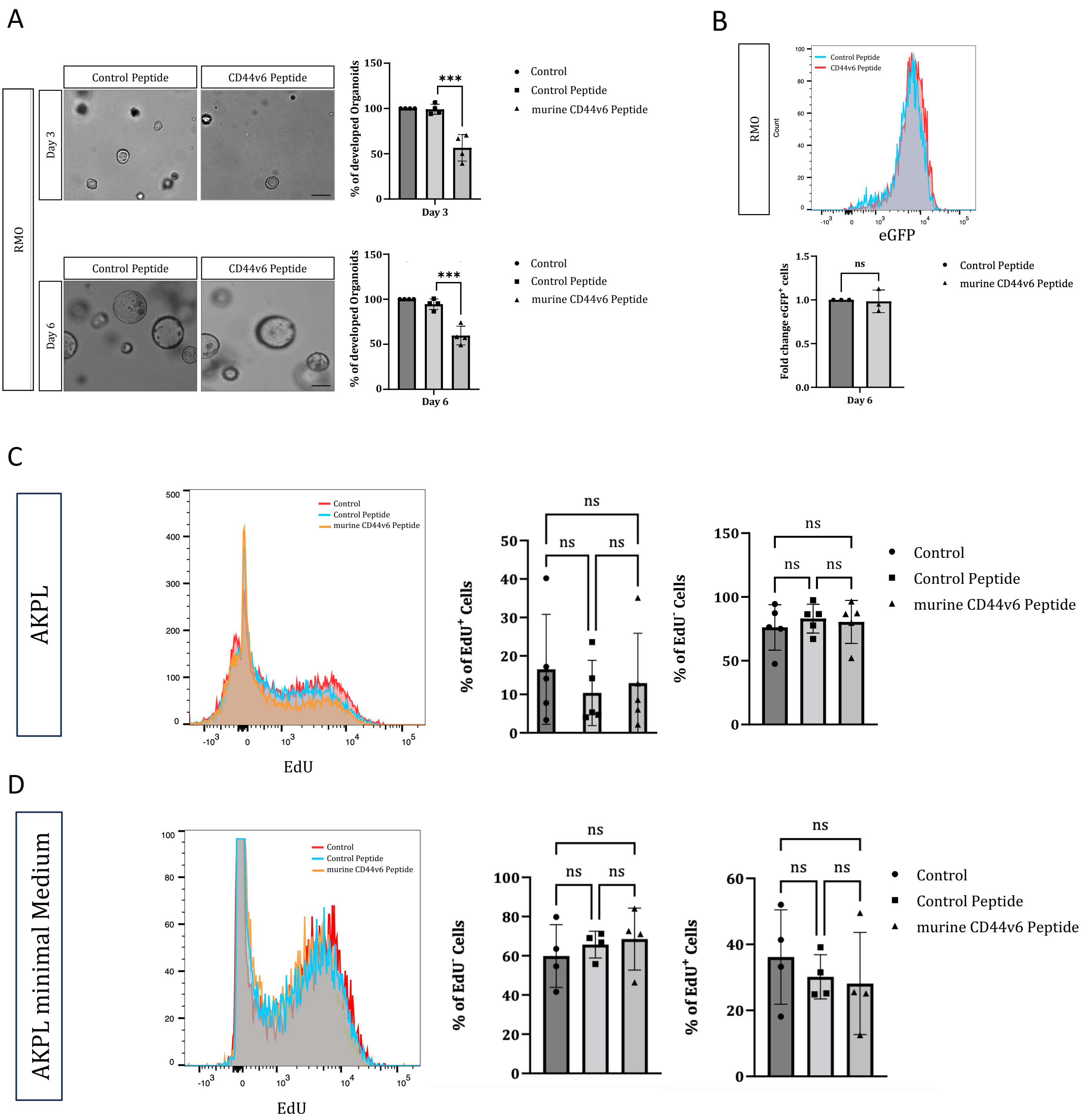

#### Supplemental Figure 4

A

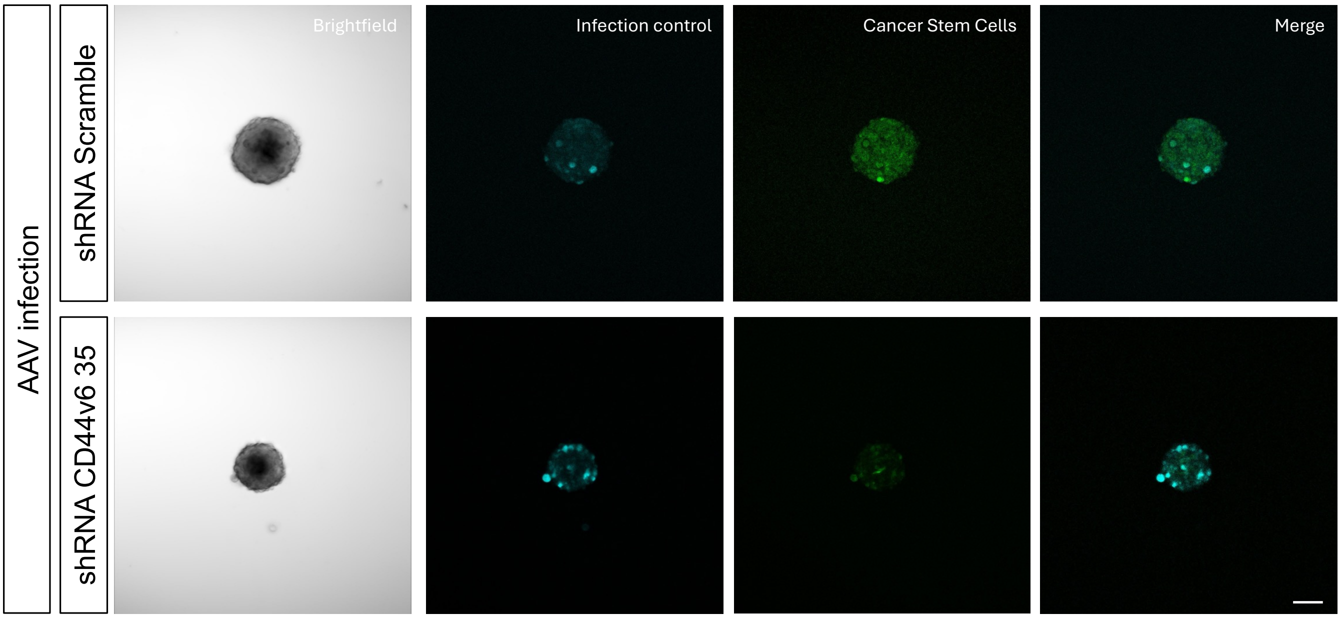


B

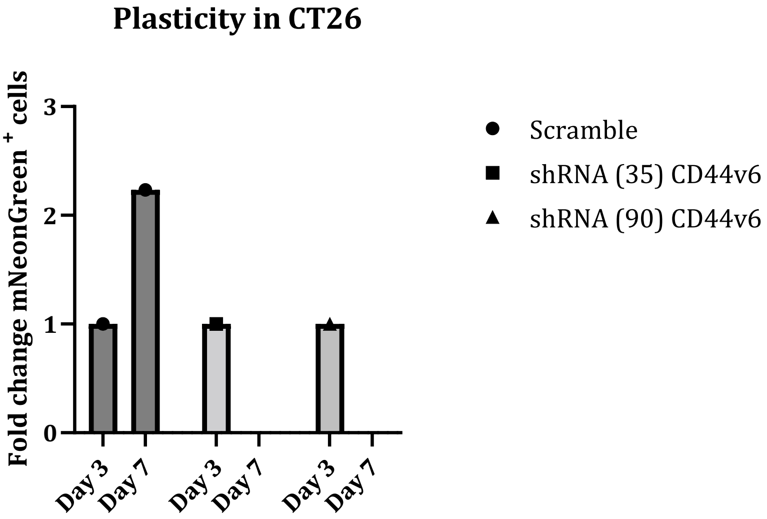
